# Benchmarking tRNA-Seq quantification approaches by realistic tRNA-Seq data simulation identifies two novel approaches with higher accuracy

**DOI:** 10.1101/2023.12.13.571582

**Authors:** Tom Smith, Mie Monti, Anne E Willis, Lajos Kalmár

## Abstract

Quantification of transfer RNA (tRNA) using illumina sequencing based tRNA-Seq is complicated due to their degree of redundancy and extensive modifications. As such, no tRNA-Seq method has become well established, while various approaches have been proposed to quantify tRNAs from sequencing reads. Here, we use realistic tRNA-Seq simulations to benchmark tRNA-Seq quantification approaches, including two novel approaches. We demonstrate that these novel approaches are consistently the most accurate, using data simulated to mimic five different tRNA-Seq methods. This simulation-based benchmarking also identifies specific shortfalls for each quantification approach and suggests that up to 13% of the variance observed between cell lines in real tRNA-Seq data could be due to systematic differences in quantification accuracy.

## Introduction

Transfer RNA (tRNA) are abundant, highly structured, short (70-100 nt) non-coding RNAs that are essential for the conversion of coding sequences into polypeptides^1^. They can be categorised by their aminoacylation identity into 20 groups, each composed of several tRNAs, known as isoacceptors, that are able to translate synonymous codons with the same amino acid. Adding to this complexity, in higher eukaryotes, there are also isodecoders: tRNAs with the same anticodon but different sequences. tRNAs also undergo substantial chemical modification, containing on average 13 modifications by molecule^2^ with over 100 post-transcriptional modifications identified to date^1,3^. Notably, a myriad of human diseases have been associated with dysregulation of tRNA expression^4–6^ and mutations in tRNA modification enzymes^7–10^, underscoring their central role in cellular homeostasis.

Given their complexity, quantification of tRNAs is more challenging than other RNA species. Experimentally, tRNAs have been intractable to high-throughput illumina sequencing due to the compact secondary and tertiary structure. In particular, the highly base-paired 5’ and 3’ termini interfere with adaptor ligation^11^, although this has been addressed in recent publications using splint adapters which take advantage of the 3’ CCA on mature tRNAs^11,12^. Moreover, some of the post-transcriptional modifications present on tRNAs disrupt the Watson-Crick face and impede reverse transcription (RT), leading to either premature termination of the RT reaction and/or high misincorporation rates around the modified nucleotide^13^. To alleviate the impact of such modification, demethylating enzymes, such as AlkB, have been utilised in several methods including DM-tRNA-Seq^14^ and ARM-seq^15^, although the activity of these enzymes have not been well characterised. An alternative solution is the use of a modification tolerant RT, such as TGIRT^14,16^, Superscript IV^12,17^ or marathonRT^18^, which have a higher processivity though modified nucleotides. Regardless of the approach taken, tRNA-Seq reads inevitably possess increased misincorporation rates and usually 3’ truncations. The processing of tRNA-Seq data is also more complex than other forms of RNA-seq, as the evolution processes exerted upon tRNA genes has led to the accumulation of dozens near identical sequences^19^. In the case of the human genome and model mammalian species such as mouse and rat, this has generated hundreds of actively transcribed tRNA genes^20,21^. This sequence redundancy complicates the easy assignment of tRNA-Seq reads to their genomic loci and, therefore, quantification is normally performed at the level of the mature tRNA sequences, or else anticodons.

Multiple quantification approaches for full length illumina-based tRNA-Seq have been employed, all relying upon alignment of reads to tRNA sequences, with considerable variability in how multi-mapped reads are handled. Bowtie2 was designed to align reads accurately and quickly^22^ and is the most frequently used aligner for tRNA-Seq^12,15,23,24^. A variety of bowtie2 parameterisations employed to alleviate the issue of misincorporation at modified bases, primarily the use of very short and/or non-exact alignment seeds, which ensures the second stage of alignment by dynamic programming has at least a minimal alignment seed from which to extend^12,23^. Aligners designed for small RNA-seq such as SHRiMP^25^ have also been used^11^. SHRiMP performs a more exhaustive search for the correct alignment and handles high error rates well, which likely makes it suitable for tRNA-Seq alignment with default parameterisation, albeit at a cost of increased runtime and memory requirements^25,26^.

Due to the high similarity between tRNA sequences, multi-mapped reads present another significant problem, which has again been solved by a variety of means, including the complete exclusion of multi-mapped reads^12,14^, fractional assignment to all aligned tRNAs^17^, and random selection from amongst the alignments^15^. Recently, mimseq, an end-to-end tRNA-Seq quantification pipeline was developed, which proposes a novel approach to deal with modified bases and multi-mapped reads^16^. This pipeline uses GSNAP^27^ to align reads to consensus sequences representing clusters of similar tRNAs along with an iterative mapping procedure, in which mutated positions are masked. Despite this significant variability between the quantification approaches which have been applied to tRNA-Seq, to date, no systematic comparison of these approaches has been performed.

Here, we used real tRNA-Seq data to generate realistic benchmark data simulations and compare common quantification approaches. This allowed us to identify the best generic quantification approach across different tRNA-Seq methods and to establish specific shortfalls of each approach.

## Results

### Optimising read alignment for tRNA-Seq

Read alignment software needs to be carefully parameterised for tRNA-Seq and a number of alternative parameterisations have been proposed, predominantly using bowtie2 with a short seed length^12,14,17^. To evaluate alignment parameters for bowtie2 with a short seed length, we simulated the highest possible quality of tRNA-Seq reads: full length and including only sequencing errors. These reads were then aligned to the reference tRNA sequences with bowtie2, using a sensitive parameterisation, allowing a single error in the seed, as frequently employed for tRNA-Seq to accommodate the increased misincorporation rate^12,23^. Two parameters were varied: L and D, representing the seed length and how many seed extensions in a row can fail to produce an improved alignment before the search is halted, respectively.

This simulation and alignment procedure was performed for tRNA from *Homo sapiens* and *Mus musculus* tRNAs separately. We observed that when an error in the seed is allowed, as the seed length is decreased, there needs to be a concomitant increase in effort expended to allow bowtie2 more opportunities to find the best possible alignment, especially with respect to the Transcript ID (Figure 1a). Thus, previously employed alignment parameters have been sub-optimal, even for unrealistically high quality tRNA-Seq reads^12^.

**Figure 1.**
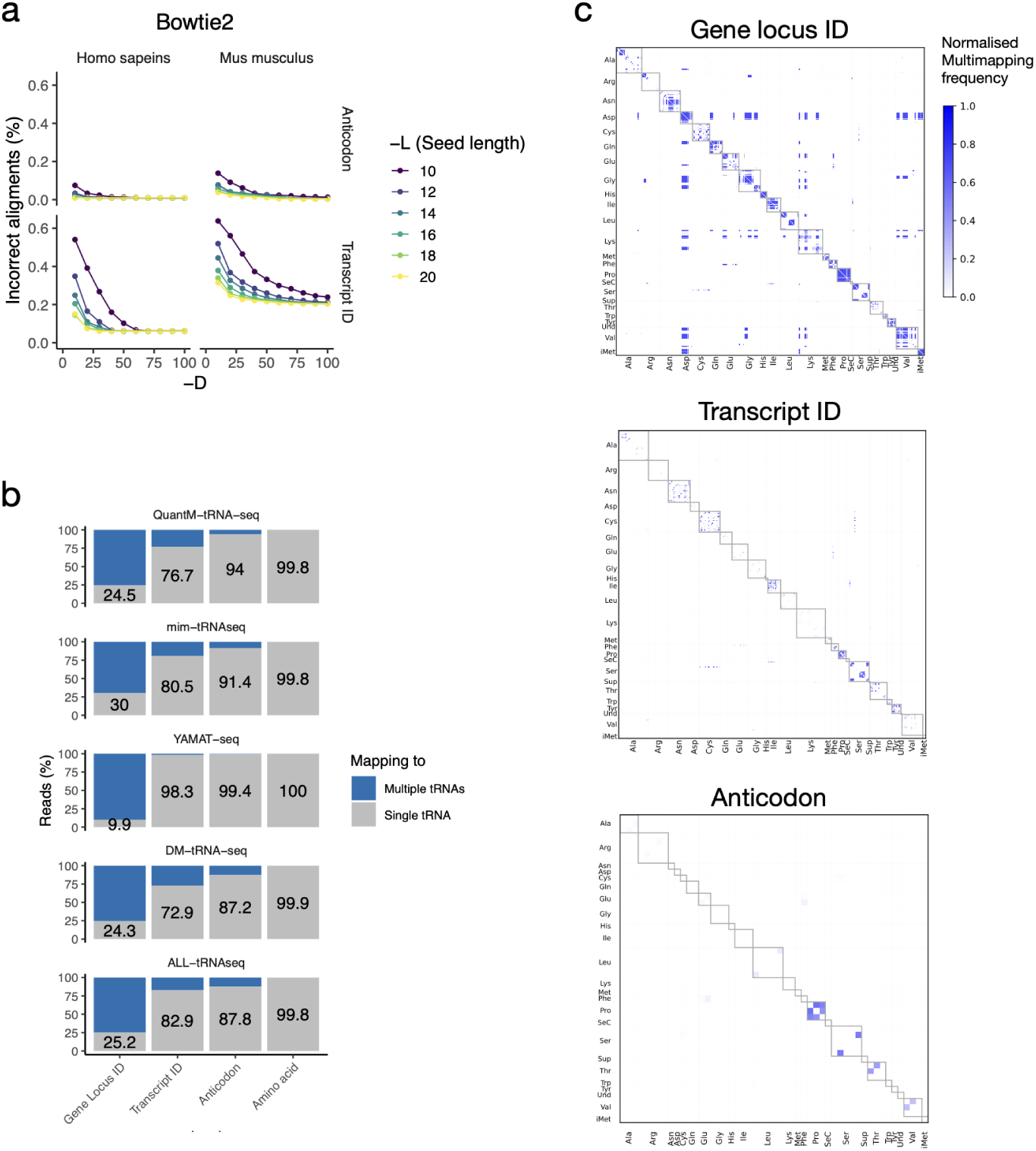
Alignment parameterisation in the context of frequent multiple-alignment. **a)**. Impact of bowtie2 parameterisation on the percentage of correct alignments. Reads were simulated as full length and with only sequencing errors. **b)** The percentage of single and multi-mapped reads at each level of tRNA nomenclature. **c).** The frequency of reads multi-mapping at the Gene locus ID, Transcript ID and Anticodon level. Frequencies were normalised by dividing by the total number of reads aligned to the two tRNAs. Data shown is from YAMAT-Seq, BT20, replicate A.

Inevitably, tRNA-Seq reads will frequently be mapped to multiple tRNA sequences with equal alignment scores, since some tRNA sequences are nearly identical and real tRNA-Seq reads will be truncated and include misincorporations. To explore the extent of the multi-mapping issue for tRNA-Seq, we aligned reads from ALL-tRNAseq^18^, DM-tRNA-seq^14^, mim-tRNAseq^16^,

QuantM-tRNA-seq^12^ and YAMAT-seq^11^ samples to the reference tRNA sequences, using bowtie2 (-L 10 -D 100). We then summarised the extent of multi-mapping at four levels using the naming convention from GtRNAdb^20^: Gene locus ID, Transcript ID, Anticodon and Amino acid.

As expected, at the higher levels of tRNA annotation, the number of reads mapping to a single feature increases (Figure 1b). Many pairs of tRNA sequences showed high rates of multi-mapping at the Gene locus ID level, including pairs of tRNAs for different anticodons, but multi-mapping is less prominent at higher levels (Figure 1c). These data demonstrate that multi-mapping is prevalent at the level of the alignments to the tRNA sequences, but it should be less problematic at higher levels if it is dealt with appropriately.

### Simulating realistic tRNA-Seq data

To address the question of how to most accurately quantify tRNAs from tRNA-Seq, we aimed to simulate realistic tRNA-Seq data by using the frequency of misincorporations and truncations (hereafter referred to jointly as the ‘error profile’) as observed in tRNA-Seq data. To ensure our findings were generically applicable, error profiles were learned from five tRNA-Seq methods: ALL-tRNAseq, DM-tRNA-seq, mim-tRNAseq, QuantM-tRNA-seq and YAMAT-seq, and two species: *Homo sapiens* and *Mus musculus*. These methods include various sequencing library preparation modifications specifically designed to handle tRNAs, primarily the use of the demethylating enzyme AlkB (ALL-tRNA-seq and DM-tRNA-seq) and/or modification tolerant RTs (ALL-tRNAseq, DM-tRNA-seq, mim-tRNAseq and QuantM-tRNA-seq), as well as the use of mature tRNA-specific double-stranded adapters (YAMAT-seq and QuantM-tRNA-seq). The error profiles obtained for each tRNA-Seq method should reflect the specific library preparation steps employed and a quantification approach which works across data simulated from all these methods should be generally applicable to tRNA-Seq. Reads were aligned to a tRNA reference transcriptome, following which, error profiles were determined from the alignments by tallying mutation frequencies and read alignment coordinates.

Since the mutations and truncations are dependent on the tRNA modifications, the position of the mutations and truncation events in the error profiles should match the positions of known modifications. Furthermore, the relationship between the tRNA modifications and error profiles should depend on whether tRNA modifications are enzymatically removed, which RT is used, and whether there was a size selection step prior to sequencing. To confirm this, we used the experimentally determined modifications listed in MODOMICS^28^. As expected, adenosines modified to inosine cause misincorporation of guanine^29^ (Figure 2a). In contrast, N1-methyladenosine (m1A) frequently causes misincorporation for all methods, except DM-tRNA-seq and ALL-tRNAseq, which employ AlkB to remove methylations. Interestingly, an overall higher proportion of mutations was observed with mim-tRNA-Seq, likely reflecting the optimisation of the RT step to allow more processivity over modified bases (Supplementary Figure 1a). In keeping with the more restrictive size selection of YAMAT-Seq, and the improved RT processivity with mim-tRNA-Seq and ALL-tRNA-Seq, the reads from these methods are less frequently truncated than the other methods (Figure 2b; Supplementary Figure 1b). Notably, while read truncations were frequently close to known modified nucleotides for all methods except YAMAT-Seq, the modified nucleotides around the site of truncation varied considerably between the tRNA-Seq methods (Figure 2c). This is likely due to the considerable differences between the tRNA-Seq library preparation steps for these five methods. Overall, the error profiles appear to reflect the different mutation and truncation rates that would be expected for the five tRNA-Seq methods and should thus enable relatively realistic simulation of tRNA-Seq reads from a broad range of tRNA-Seq protocols.

**Figure 2.**
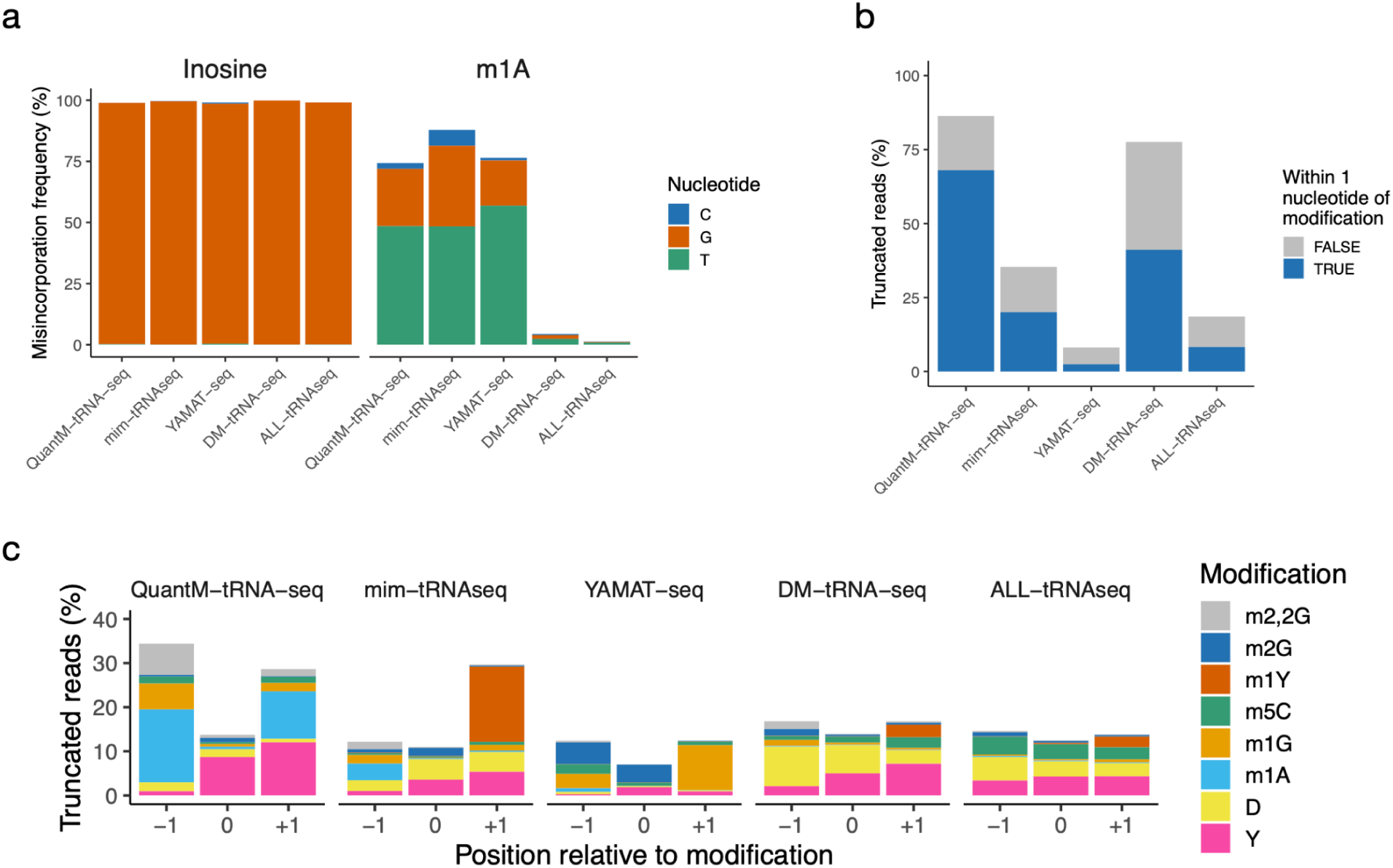
The observed error profiles reflect the expected misincorporations from modifications in MODOMICS. **a)**. Misincorporation frequencies for Inosine and m1A sites **b)**. The proportion of reads which are truncated and how often this occurs close to a known modification site. **c)**. The proportion of truncated reads which are truncated within 1 nucleotide of common modifications. m2,2G = N2,N2-dimethylguanosine, m2G = N2-methylguanosine, m1Y = 1-methylpseudouridine, m5C = 5-methylcytidine, m1G = 1-methylguanosine, m1A = 1-methyladenosine, D = dihydrouridine, Y = pseudouridine.

From these error profiles, we then simulated 2 datasets: *Uniform*: an equal number of reads from all tRNA species, with misincorporation and truncations added at the frequencies observed in the real tRNA-Seq data. One set of simulated reads was generated for each tRNA-Seq sample. *Realistic*: a randomly varied number of reads from each tRNA sequence, drawn from a log2-Gaussian distribution with a mean of 10 and standard deviation of 5, left-censored at zero. Misincorporation and truncations were added at the frequencies observed in the real tRNA-Seq data. Ten sets of simulated reads were generated for each tRNA-Seq sample.

### Comparing alignment strategies

The first question we wished to address was which read alignment strategy yielded the most accurate alignments. We therefore compared 3 strategies using the *Uniform* simulation data by employing either bowtie2 or SHRiMP alignment to tRNA sequences or alignment to consensus tRNA sequences using GSNAP, using the mimseq pipeline^16^. Bowtie2 and SHRiMP alignments were analysed at 3 levels using the naming convention from GtRNAdb: Gene locus ID, Transcript ID and Anticodon. The mimseq pipeline aligns to consensus tRNA sequences that approximate a single transcript ID and are associated with a single anticodon. Therefore, we also analysed the bowtie2 and SHRiMP alignments at the ‘Mimseq isodecoder’ level to make them comparable.

With the configurations used here, bowtie2 and SHRiMP aligned the most reads, with the median percentage of aligned reads across all *Homo sapiens* samples being 98.5%, 93.0% and 71.3%, for bowtie2, SHRiMP and mimseq (GSNAP), respectively (Figure 3a). However, relative to mimseq, SHRiMP and bowtie2 had up to 3.3% and 4.0%, respectively, fewer alignments correct at the anticodon level for *Homo sapiens* samples (Figure 3b; supplementary Figure 2a). This suggests there is an expected trade-off between the proportion of aligned reads and the accuracy of the alignments (Supplementary Figure 2b).

**Figure 3.**
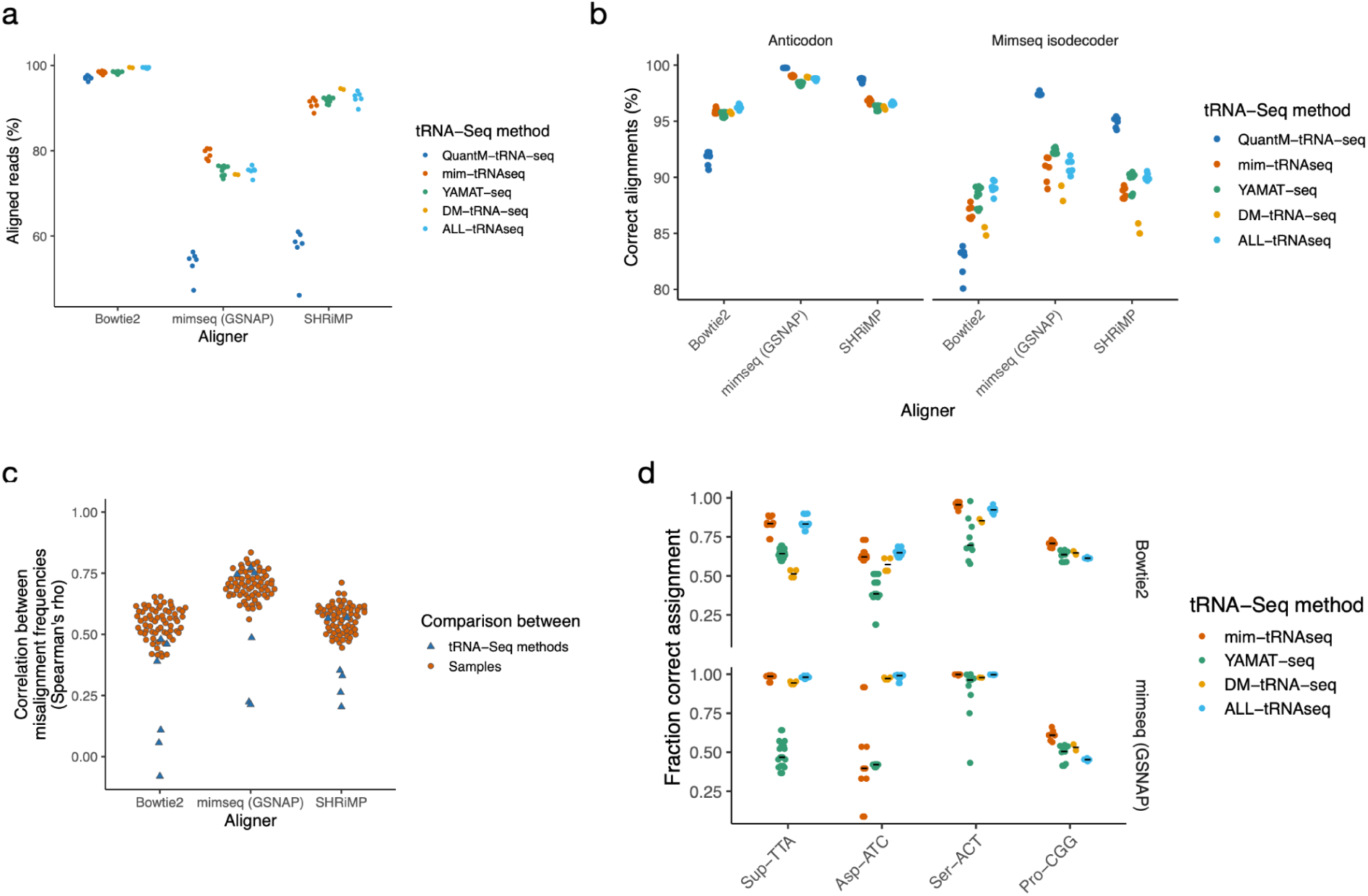
Alignment accuracy is dependent upon the alignment strategy and tRNA-Seq method. **a)** Percentage of reads aligned. **b)**. Percentage of correct alignments. **c)**. Spearman correlations for read misalignments rates between pairs of tRNA-Seq methods or pairs of samples within a single tRNA-Seq method. QuantM-tRNA-seq is not included since these samples were only for *Mus musculus*. **d)**. A comparison of correct assignments for 4 selected *Homo sapiens* anticodons which are most variable across tRNA-Seq methods. QuantM-tRNA-seq is not included since these samples were only for *Mus musculus*.

We next considered the consistency for the read misalignments patterns. Notably, the read misalignments were more correlated between samples from the same tRNA-Seq method than between the two tRNA-Seq methods for each species (Figure 3c), likely due to the differences in error profiles between each tRNA-Seq method (Figure 1). Interestingly, some tRNAs anticodons were difficult to align when simulating reads from specific tRNA-Seq methods. For example, using bowtie2, reads from *Homo sapiens* Sup-TTA tRNAs are more correctly aligned for YAMAT-seq than DM-tRNA-seq. In contrast, mimseq accurately aligned Sup-TTA reads simulated from DM-tRNA-seq, but not from YAMAT-seq (Figure 3d).

We also observed cell-line specific misalignment patterns that vary across the tRNA-Seq methods and alignment approaches (Supplementary Figure 2c). For example, using mimseq, reads simulated from mim-tRNAseq for Asp-ATC tRNAs were missassigned to Cys-GCA tRNAs 67%, 40% and 100% of the time for BT20, MCF7, and SKBR3, respectively. However, using bowtie2, the misalignment rates were only 18%, 24% and 36%, respectively. The difference in the misalignments between the cell lines and tRNA-Seq methods supports the approach taken herein to use learn sample-specific error profiles from multiple tRNA-Seq methods to enable simulation of a broad set of representative samples from which to identify a generically accurate quantification approach.

### Handling multiple mapped reads

Once reads are aligned, the major variable is how to handle multi-mapped reads in the process of tallying read counts per tRNA. To compare approaches, we aligned the *Realistic* simulated reads to tRNA sequences with bowtie2 or SHRiMP and applied 4 approaches used previously to count reads per tRNA, alongside two further approaches which we expected would be more accurate that have not previously been applied, to our knowledge (Table 1).The Decision approach only assigns reads to a tRNA if they are uniquely mapped, but re-frames the uniqueness of the mapping with respect to the level of tRNA quantification being performed. For example, a read aligning to multiple tRNA sequences with the same transcript ID would be assigned to that transcript ID and the associated Anticodon, but would be discarded for Gene locus ID quantification. Salmon implements a lightweight mapping procedure and a maximum-likelihood approach to estimate RNA abundances from the mapped reads, while accommodating multi-mapped reads^30^. To ensure the quantification approaches were comparable, we used Salmon in the ‘alignment-based’ mode, and provided the same read alignments from bowtie2 or SHRiMP which were used for the other read tallying approaches. In addition, we used the dedicated tRNA-Seq quantification tool, mimseq, which performs a bespoke alignment and quantification.

**Table 1.**
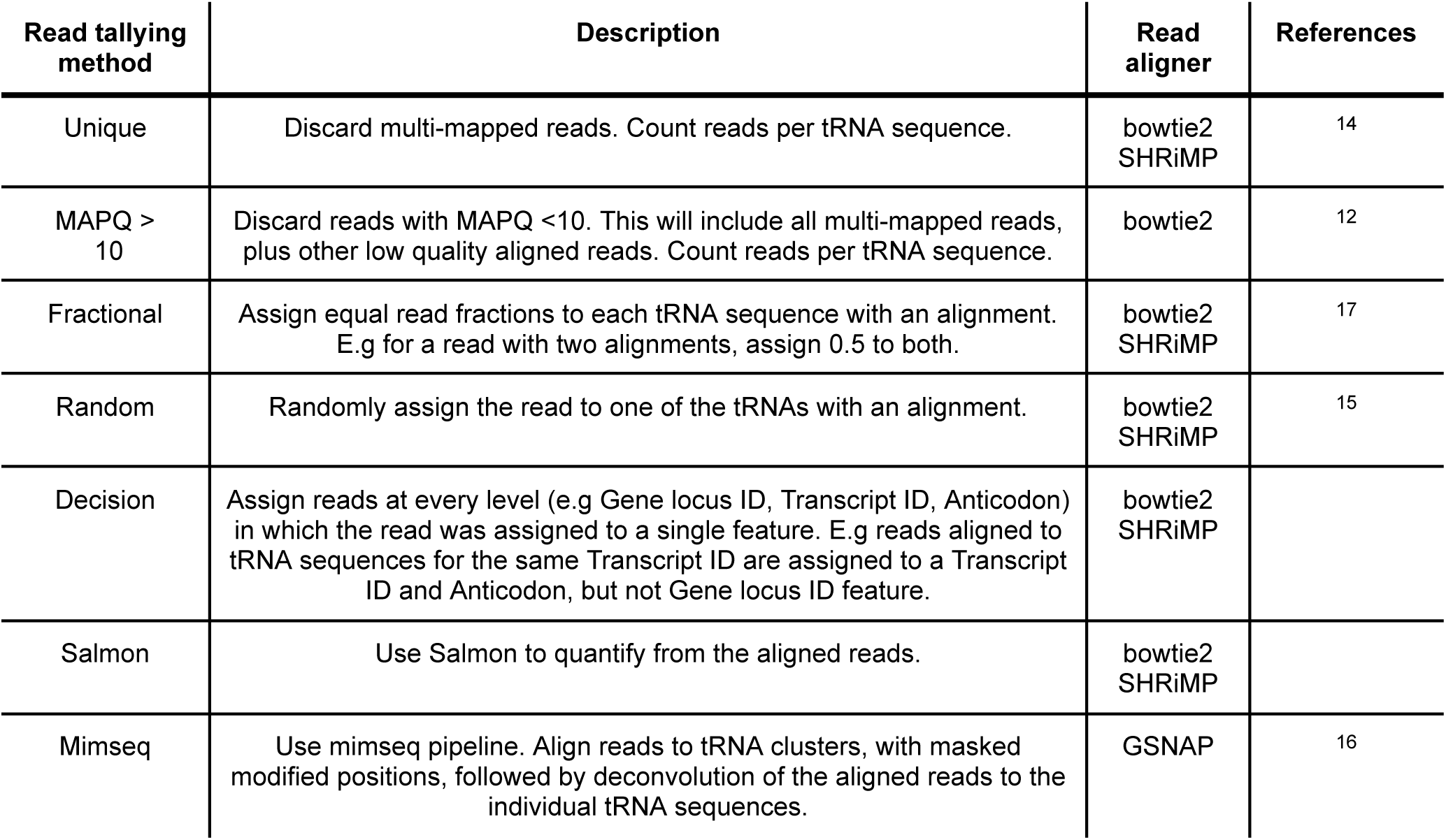
Approaches for tallying reads per tRNA from aligned reads. SHRiMP does not report MAPQ, so the ≠=MAPQ>10 approach is not possible. Decision and Salmon have not previously been utilised.

| Read tallying method | Description | Read aligner | References |
| --- | --- | --- | --- |
| Unique | Discard multi-mapped reads. Count reads per tRNA sequence. | bowtie2<br>SHRiMP | 14 |
| MAPQ > 10 | Discard reads with MAPQ <10. This will include all multi-mapped reads, plus other low quality aligned reads. Count reads per tRNA sequence. | bowtie2 | 12 |
| Fractional | Assign equal read fractions to each tRNA sequence with an alignment. E.g for a read with two alignments, assign 0.5 to both. | bowtie2<br>SHRiMP | 17 |
| Random | Randomly assign the read to one of the tRNAs with an alignment. | bowtie2<br>SHRiMP | 15 |
| Decision | Assign reads at every level (e.g Gene locus ID, Transcript ID, Anticodon) in which the read was assigned to a single feature. E.g reads aligned to tRNA sequences for the same Transcript ID are assigned to a Transcript ID and Anticodon, but not Gene locus ID feature. | bowtie2<br>SHRiMP |  |
| Salmon | Use Salmon to quantify from the aligned reads. | bowtie2<br>SHRiMP |  |
| Mimseq | Use mimseq pipeline. Align reads to tRNA clusters, with masked modified positions, followed by deconvolution of the aligned reads to the individual tRNA sequences. | GSNAP | 16 |

With the exception of Decision and mimseq, which inherently generate quantification at multiple levels, all other approaches quantify at the level of the Gene locus ID sequences included in the reference fasta. We further summarised quantification to transcript ID and anticodon-level feature abundances by summation. We also summarised the quantification values to the same ‘mimseq isodecoder’ features quantified by mimseq, to make them comparable. A summary of the 4 levels of quantification and how these are obtained for each read tallying approach is shown (Table 2).

**Table 2.** Quantification levels, using GtRNAdb nomenclature, and how quantification is achieved at each level by each read tallying approach.

| Quantification feature level | Description | Obtained directly | Obtained by summation |
| --- | --- | --- | --- |
| Gene locus ID | Unique tRNA genomic loci | Unique, MAPQ > 10<br>Fractional, Random,<br>Decision, Salmon, Mimseq |  |
| Transcript ID | tRNAs with the same mature sequence | Decision | Unique, MAPQ > 10<br>Fractional, Random<br>Salmon |
| Mimseq isodecoder | Clusters of similar sequences defined by mimseq pipeline in data-dependent manner<sup>16</sup> | Mimseq | Decision, Salmon, Unique,<br>MAPQ > 10, Fractional,<br>Random, Salmon |
| Anticodon | tRNAs with the same anticodon | Mimseq, Decision | Unique, MAPQ > 10,<br>Fractional, Random, Salmon |

The proportion of reads used by each read tallying approach varies considerably. *Salmon*, *Fractional* and *Random* use all mapped reads for each level of quantification, whereas stricter approaches which filter by multimapping or mapping quality use the least reads and frequently assign fewer than 50% of the mapped reads (Figure 4a; Supplementary Figure 3a). As intended, *Decision* uses increasingly more reads at higher levels of quantification. Importantly, *Decision* uses approximately the same number of reads as mimseq at the mimseq isodecoder level, but consistently more at the anticodon level, where it approaches the use of all mapped reads. In contrast, *Mimseq* uses the same number of reads for both mimseq isodecoder and anticodon level quantification. This suggests *Decision* is more appropriately determining which reads to utilise for each level of quantification.

**Figure 4.**
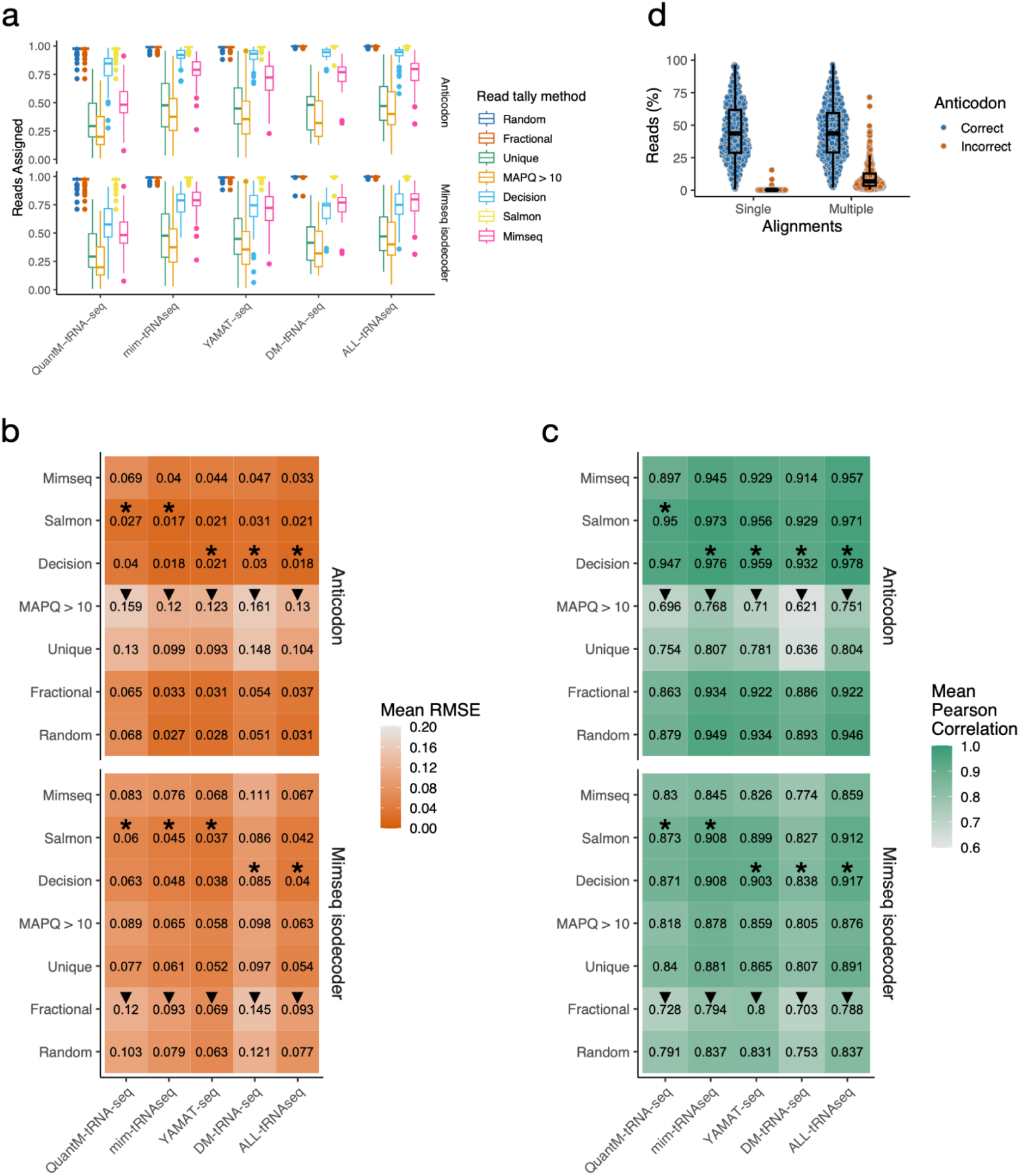
**a)** Reads assigned at anticodon and mimseq isodecoder level for each read tally method. Only results from alignment with bowtie2 or GSNAP (Mimseq) are shown. **b)** Mean RMSE at anticodon and mimseq isodecoder level. *=Best approach for each tRNA-Seq method, ▾=Worst approach. **c)**. As per (b), except Pearson correlation coefficient. **d)** The percentage of reads from each simulation sample which are single or multiple aligned, separated by whether the alignment position(s) have the correct anticodon. Multi-aligned reads with more than more anticodon were deemed incorrect.

### Comparing read tallying approaches

To assess the overall accuracy of the abundance estimates obtained with the varying read tallying approaches, we calculated two metrics. The root mean squared error (RMSE) is a common error metric which captures how closely the quantification estimates match the ground truth^31^. However, in many applications of tRNA-Seq, including the comparison between conditions or tissues, it’s sufficient to accurately capture the abundance relationship between samples, even if there is a systematic over or under-estimation across all samples. Hence, we complemented the evaluation using RMSE by also considering the Pearson correlation coefficient between the quantification estimates and ground truth.

To compare the quantification approaches, we considered the mean RMSE and mean correlation across all features at a given quantification level. At the anticodon and mimseq isodecoder level, our novel approaches, *Decision* and *Salmon* consistently show the lowest RMSE and highest correlation, suggesting they deal with multi-mapped reads most appropriately. Of the previously used read-tallying approaches, *Mimseq* shows the highest accuracy. Surprisingly, *Random* and *Fractional* also perform well at the anticodon level, while *Unique* and *MAPQ>10* performed poorly (Figure 4b-c). On further inspection, this is because the multi-mapped reads are predominantly mapped to multiple tRNA species from the same anticodon (Figure 4d; Supplementary Figure 3a). Thus, requiring unique alignments at the level of individual tRNA sequences (using *Unique* or *MAPQ>10)* leads to detrimental discarding of multi-mapped reads which could be assigned to a single anticodon. For these reads, selecting one alignment at random, or equally distributing the read between all of them, is a more sensible approach and thus leads to more accurate anticodon level quantification. However, for quantification at lower levels, Unique and *MAPQ > 10* are more accurate than *Random* and *Fractional*, suggesting discarding of multi-mapped reads is better than naive resolution for quantification below the anticodon level (Figure 4b-c). Very similar results are observed when SHRiMP is used in place of bowtie2 (Supplementary Figure 3b-c). Finally, for quantification at the level of individual tRNA sequences, all methods perform equally well, with the exception of *Random*, which performed the worst. Overall, *Decision* and *Salmon* are consistently the best read tallying methods to apply.

We reasoned that the misalignment of reads should be one of the major factors reducing quantification accuracy for all methods. In support of this, the fraction of reads correctly aligned and the quantification evaluation metrics were correlated (Supplementary Figure 4a-b). The exception was *MAPQ>10* at anticodon level, which shows poor accuracy even when the proportion of correctly aligned reads is high. This agrees with the previous observation that discarding low quality reads is detrimental for anticodon-level read tallying because reads are typically mapped across multiple tRNAs for the same anticodon, not between anticodons (Figure 3d). Thus, excepting quantification by *MAPQ>10,* our results indicate that misalignment is one of the major issues with tRNA-Seq quantification.

### Using simulations to inform interpretation of results

Summarising the RMSE and correlation metrics to the mean value over all features enables read tallying approaches to be easily compared, but masks fine-grained differences which go against the trend and could give false confidence when interpreting quantification of individual features. We therefore further considered which features each approach quantifies best, focusing on the simulations from the YAMAT-Seq data and quantification at the anticodon level. For these simulations, *Decision* had the highest mean correlation and lowest mean RMSE, with *Salmon* performing nearly equivalently and *Mimseq* also being accurate (Figure 4c). However, we observed that the correlations for some anticodons were very sensitive to the read tallying approach (Figure 5a-b). For example, the correlation for Ser-GGA is much lower with *Mimseq* than *Decision* or *Salmon*, whilst the correlation for Pro-AGG is much higher with *Mimseq* than *Decision* or *Salmon* (Figure 5b; Supplementary Figure 4c). This suggests the choice of quantification approach may have different impacts on quantification accuracy for each anticodon and the interpretation of results should ideally take into account the limits of each quantification approach.

**Figure 5.**
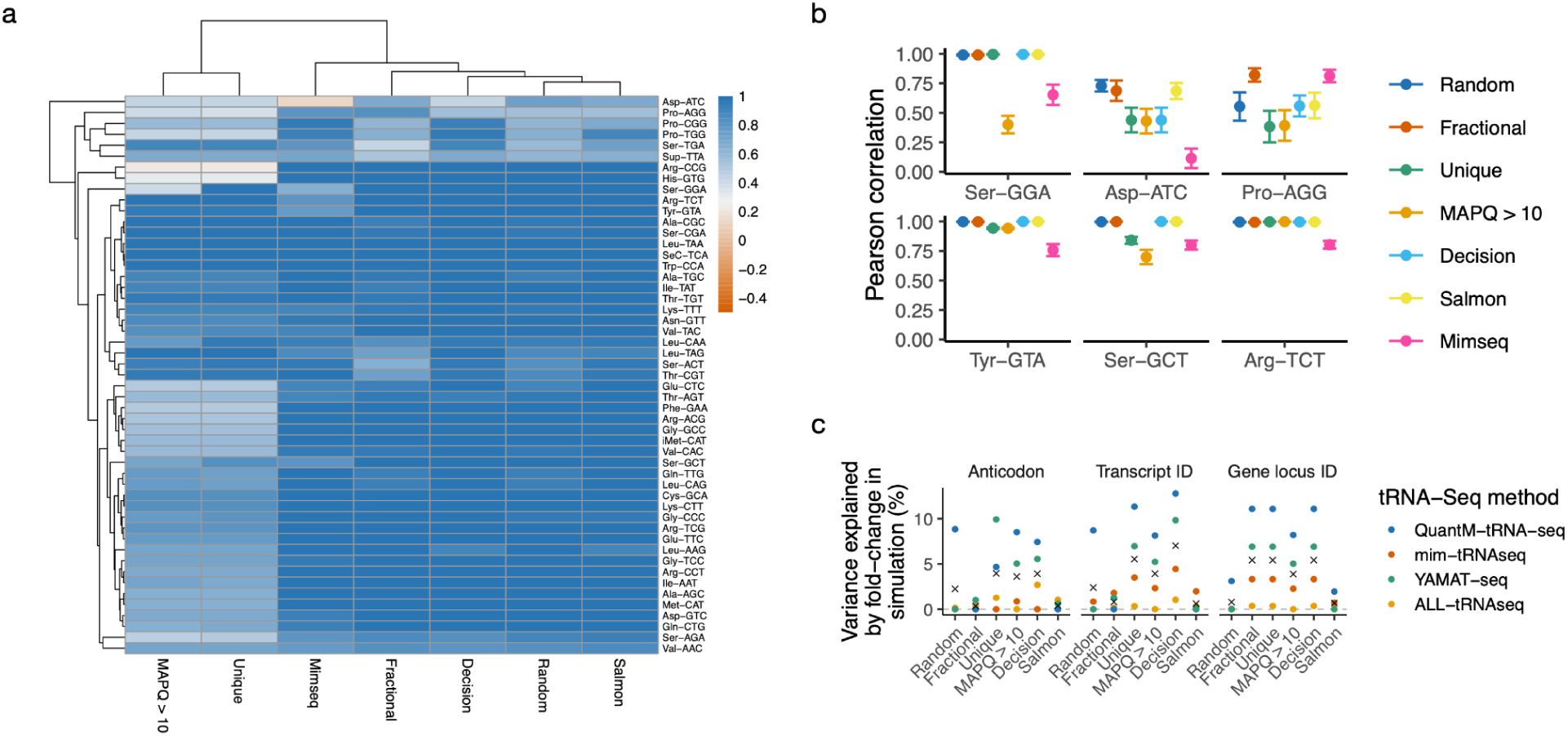
Each quantification approach has specific shortfalls. **a)** Pearson correlation between ground truth and estimated read counts for six anticodons with variable quantification accuracy between read tallying approaches. **b). c)** The variance in observed fold-changes in the real tRNA-Seq data that is explained by fold-changes in the uniform simulation data. x = The mean variance explained across the four *Homo sapiens* datasets.

Given the observed association between read misassignment and quantification accuracy, we hypothesised that differences in read misalignments between cell lines could erroneously induce apparent differences in tRNA feature abundances. To test this, we compared the fold-change between cell lines when simulating the same number of reads for each tRNA with the fold-changes observed in the real data. The fold-changes with the simulated data should reflect technical biases resulting from systematic differences in misassignments between cell lines due to differences in the error profiles. Strikingly, up to 13% of the variance in fold-changes in the real data were explained by technical biases (Figure 5c). Even using the most accurate quantification method, *Decision*, up to 7% of the variance for anticodon level fold-changes in the real data is explained by technical biases. All quantification methods appeared to show some propensity to identify fold-changes between conditions that were partially explained by technical biases for at least one level of tRNA quantification. The exception to this was salmon, for which a maximum of 2% of the variance in fold-changes were explained by technical biases. This suggests salmon may be a more trustable quantification method for the detection of fold-changes between experimental conditions.

## Discussion

Recent advances in sequencing using nanopores promise a potential revolution for RNA sequencing, including tRNAs^32^. Despite this, illumina sequencing is expected to remain the dominant approach for tRNA sequencing, at least in the short-term. The first tRNA-seq approaches using illumina sequencing were published in 2015^14,15^, since which, the comparison of bioinformatic approaches for tRNA quantification from tRNA-Seq has received surprisingly little attention. In part, this may be because tRNA-Seq methods continue to be developed and the optimal quantification is considered to be dependent upon the method. Encouragingly, however, we have identified two robust quantification methods that appear to be suitable for all tRNA-Seq data.

An important aspect of our benchmarking is the simulation of realistic tRNA-Seq data, which has not been attempted before, to our knowledge. The only previous attempt to simulate tRNA-Seq reads that we are aware of did not capture the effect of modifications on read truncations and mutations^33^. By learning error profiles of individual samples and then simulating data from these error profiles, our simulations at least partially capture the technical variability between samples, cell lines and tRNA-Seq methods. However, there are limitations to this approach. All truncation and misincorporation events are measured independently, such that the error profiles will not capture dependencies between the events, which occur with respect to correlated misincorporations at nucleotides due to linked modifications^34^. In addition, the error profiles are inherently biased by the ability to align the tRNA-Seq reads. Misincorporations and truncations which prevent the read being aligned appropriately will not be captured in the error profiles. As such, the simulated data may be considered to simulate the alignable portion of a tRNA-Seq sample, with biases due to the alignment strategy used. Here, we attempted to mitigate this by aligning reads with multiple aligners. Despite these limitations, our simulations represent the most realistic simulation of tRNA-Seq data to date, and enabled us to perform a rigorous comparison of tRNA-Seq quantification approaches.

The crucial steps for accurate tRNA-Seq quantification are the alignment of reads and handling of multi-mapped reads, with the latter being the step which varies most between the approaches that have been applied to date. Although reads may be multi-mapped across different tRNA species, we show here that the issue is most pressing at the Gene locus ID and Transcript ID level, and much less so at the Anticodon level. However, dealing with multi-mapped reads inappropriately can itself create issues at the anticodon level. The approaches used to date resolve the multi-mapping are only suitable for quantification at particular levels, which has not previously been appreciated. Crucially, the common approach of removing multi-mapped reads and/or low quality reads is significantly suboptimal for the anticodon level quantification which it has been previously employed for^12,14^, since it leads to the unnecessary discarding of reads which have multi-mapped across tRNAs with the same anticodon. The bespoke quantification approach of *Mimseq* is better than filtering multi-mapped reads, but our simulations indicate it performed no better than simple random assignment of multi-mapped reads for anticodon-level quantification.

The central observation of this benchmarking study is that two simple approaches alleviate the issue of multi-mapped reads and perform better than existing approaches at all levels of quantification. This is observed across all 4 of the tRNA-Seq methods, suggesting our observation can likely be extrapolated to all tRNA-Seq methods using illumina sequencing for full length tRNAs. The *Salmon*-based approach involves reporting all alignment positions and then using salmon to probabilistically resolve the multi-mapped reads. The *Decision* approach involves counting reads at each level of quantification where the read maps to tRNAs from a single feature, e.g multiple tRNAs from the same anticodon. Given the lower propensity for technical biases to explain fold-changes in real data with *Salmon* and the reduced barrier to its use, we suspect salmon will prove to be the most popular. For this approach, bowtie2 simply needs to be run with the -a flag to report all reads. The alignments must then be filtered to retain only the equal top-scoring alignments for each read, before passing the alignments to salmon quant.

While the motivation of this study was to determine the optimal overall quantification approach, our simulations also identified specific tRNAs that cannot be accurately quantified in particular samples. Our simulations further allowed us to establish that a small proportion of the observed variability between cell lines may not be biological. Thus, comparisons between samples should be interpreted with caution, since they may be partially due to technical biases in read miss-alignment. We propose that a simulation-based approach could be applied more widely as a quality control step to highlight features whose quantification may be inaccurate. In particular, where changes in tRNA abundance between cell lines are observed, it would be prudent to use similar simulations to demonstrate that such changes are not driven by differences in tRNA modifications driving systematic differences in misincorporations and truncations.

The decision about what level to interrogate tRNA abundance is a trade off between a desire to obtain the most ‘fine-grained’ data possible, and the practical consideration of how accurately tRNAs can be quantified at each level. tRNA isodecoders have been found to be functionality distinct^35,36^ and differential abundance of specific Transcript IDs has been identified across mouse tissues^12^ and human cell lines^16^. Nonetheless, tRNA abundances are more accurately quantified at the Anticodon level and this is likely to remain the predominant approach for the foreseeable future. As tRNA-Seq methods continue to improve the processivity of reverse transcriptase over modified bases and yield fewer truncated reads, the potential for highly accurate Transcript ID level quantification will open up. Sequencing tRNAs using nanopores is also likely to provide a significant benefit in this respect^32^. However, these improvements in experimental methods need to be complemented by rigorous improvements in the data processing to take full advantage of the potential benefits. We envisage the simulation-based approach we have used here being reutilised as new experimental methods are developed, to maintain a high standard of tRNA-Seq data analysis.

## Data and code availability

The data processing pipeline is available at https://github.com/MRCToxBioinformatics/pipeline_compare_trnaseq_quant and the version used (v0.1) is archived at zenodo, DOI: 10.5281/zenodo.10229154. The pipeline makes extensive use of a dedicated python package we developed for the simulation of tRNA-Seq samples, simulatetrna, which is available at https://github.com/MRCToxBioinformatics/simulate_trna and the version used (v0.1) is archived at zenodo: DOI: 10.5281/zenodo.10235438. The downstream data analysis and visualisation was performed using R markdown notebooks ^37^, which are available at https://github.com/MRCToxBioinformatics/trna_seq_quantification_benchmarking (v0.1.1), archived with zenodo, DOI: 10.5281/zenodo.10372994.

## Acknowledgements

The original motivation to benchmark tRNA-Seq quantification approaches came from discussions with Tuija Poyry and Thomas Mulroney and we would like to thank them for their continued insightful comments and suggestions. TS, LS, MM and AEW are supported by the Medical Research Council, grant number RG94251

## Author contributions

T.S: Conceptualization, Data Curation, Formal Analysis, Methodology, Project administration, Investigation, Software, Visualization, Writing - Original Draft, Writing - Review & Editing. MM: Investigation, Visualization, Writing - Original Draft, Writing - Review & Editing. AEW: Project administration, Funding acquisition, Writing - Review & Editing. LK: Project administration, Supervision, Writing - Review & Editing.

## Copyright

For the purpose of open access, the author has applied a Creative Commons Attribution (CC BY) licence to any Author Accepted Manuscript version arising from this submission.

## Methods

A CGAT-core ^38^ python pipeline was used to generate error profiles, compare errors with known modifications, simulate tRNA-Seq samples, align reads, run mimseq, compare simulation ground truth and observed alignments, tally tRNA counts and compare simulation ground truths and quantification estimation, as described below. The pipeline is available from https://github.com/MRCToxBioinformatics/pipeline_compare_trnaseq_quant and the version used (v0.1) is archived at zenodo, DOI: 10.5281/zenodo.10229154.

Downstream data analysis and visualisation was performed on a macOS (12.1×86_64-apple-darwin17) with R v4.1.2, using Rstudio v2023.09.0+463. The Rmarkdown notebooks are available from https://github.com/MRCToxBioinformatics/trna_seq_quantification_benchmarking (v0.1.1), archived with zenodo, 10.5281/zenodo.10372994.

### Software versions

All data-processing software was installed in a dedicated conda environment. A yaml file detailing the software versions is included in the github repository for the data-processing pipeline, https://github.com/MRCToxBioinformatics/pipeline_compare_trnaseq_quant (v0.1), archived at zenodo, DOI: 10.5281/zenodo.10229154. Downstream data analysis and visualisation was performed with R v4.1.2. R package dependencies and the versions used are detailed in the github repository for the data analysis and visualisation, https://github.com/MRCToxBioinformatics/trna_seq_quantification_benchmarking (v0.1.1), archived with zenodo, DOI: 10.5281/zenodo.10372994.

### Data acquisition

*Homo sapiens* mim-tRNAseq samples were downloaded from https://www.ebi.ac.uk/ena/browser/view/PRJNA639839. These comprised two replicates each for iPSC, HEK293T and K562. *Homo sapiens* YAMAT-Seq data were downloaded from https://www.ebi.ac.uk/ena/browser/view/PRJNA360886. These comprised 3 replicates each for BT-20, SK-BR-3 and MCF-7. *Homo sapiens* DM-tRNA-Seq were downloaded from https://www.ebi.ac.uk/ena/browser/view/PRJNA277309. These comprised 2 replicates of HEK293T. *Homo sapiens* ALL-tRNASeq were downloaded from https://www.ebi.ac.uk/ena/browser/view/PRJNA775872. These comprised 3 replicates of hESC at day 0 and day 5 of retinoic acid differentiation. *Mus musculus* QuantM-tRNA-Seq data were downloaded from https://www.ebi.ac.uk/ena/browser/view/PRJNA593498. These comprised 3 replicates each from Cortex and Cerebellum. Mature tRNA-Seq sequences, mitochondrial tRNA sequences and tRNAscan-SE output files were obtained from v1.3.8 of the mimseq GitHub repository (https://github.com/nedialkova-lab/mim-tRNAseq/releases/tag/v1.3.8). Copies of the tRNA sequence files used are contained in the GitHub repository for the data processing pipeline: https://github.com/MRCToxBioinformatics/pipeline_compare_trnaseq_quant; version used (v0.1) archived with zenodo: DOI: 10.5281/zenodo.10229154

### Measuring multi-mapping

Reads from YAMAT-Seq, DM-tRNA-Seq, ALL-tRNASeq, QuantM-tRNA-Seq and mim-tRNAseq were aligned to the tRNA sequences with bowtie2^22^. bowtie2 was run with the following non-default parameters: --min-score G,1,8 --local -a -D 20 -R 3 -N 1 -L 10 -i S,1,0.5, where -a makes bowtie2 report all read alignments. The bam.filter_sam function from simulatetrna was used to filter read alignment BAM files from bowtie2 and SHRiMP to remove alignments whose alignment score was less than the highest alignment score for the read. To summarise the multimapping for each sample, the number of reads uniquely aligning to each tRNA sequence and multimapping between each pair of tRNA sequences was computed. The number of multimapping reads for each tRNA sequence pair was normalised to 0-1 through dividing by the sum of the multimapped reads and the uniquely mapped reads for each of the tRNA sequences. This procedure was performed at all levels of tRNA sequence nomenclature, as described by GtRNAdb^20^. To achieve this, the tRNA sequences to which a read was aligned (Gene locus ID level) were summarised to the higher levels of nomenclature (e.g Anticodon level) by taking the set of unique tRNAs when expressed at the higher level. For example, a read multimapping to the tRNA sequences for Val-CAC-8-1 and Val-CAC-9-1 (Gene locus ID level) would be multi-mapped to Val-CAC-8 and Val-CAC-9 at the Transcript ID level and uniquely mapped to Val-CAC at the anticodon level. Thus, reads multimapping at lower levels could be deemed uniquely mapping at higher levels.

### Obtaining error profiles

Error profiles for each sample were obtained by aligning reads to the reference tRNA sequences using bowtie2^22^ and BWA-MEM^39^, reporting a maximum of one alignment per read. Bowtie2 was run with the following non-default parameters: --min-score G,1,8 --local -D 100 -R 3 -N 1 -L 10 -i S,1,0.5. BWA-MEM was run with the following non-default parameters: -k 10 -T 15. The bowtie2 and BWA-MEM alignments for each sample were merged and passed to alignmentSummary.clustalwtrnaAlignmentSummary from the simulatetrna python library to obtain the error profiles. Error profiles represent the frequency of misincorporations at each position in each tRNA, the frequency for each observed read start site (3’ with respect to tRNA), and the frequency of truncations (5’ with respect to tRNA sequence) for each observed read start site. The error profiles do not encode dependencies between misincorporations at each position and between misincorporations and truncations. To avoid error profiles for rare tRNA sequences being inaccurately measured, error profiles were also summarised within tRNAs sharing the same anticodon for the purpose of simulating tRNA-Seq reads.

#### Comparing error profiles to known modifications

Modification data was obtained from MODOMICS on 23 January 2023 using their API, with the URLs https://www.genesilico.pl/modomics/api/sequences?RNAtype=tRNA&organism=Homo+sapiens&format=json and https://www.genesilico.pl/modomics/api/sequences?RNAtype=tRNA&organism=Mus+musculus&format=json. The details of MODOMICS modifications were downloaded from https://genesilico.pl/modomics/modifications on 23 January 2023. Copies of the MODOMICS files used are contained in the GitHub repository for the data processing pipeline: https://github.com/MRCToxBioinformatics/pipeline_compare_trnaseq_quant; version used (v0.1) archived with zenodo, DOI: 10.5281/zenodo.10229154.

MODOMICS defines modifications for only a minor portion of tRNAs. On the assumption that tRNAs which share the same anticodon and have a very similar sequence to the MODOMICS tRNA sequence will likely have the same modifications, the positions of the modifications in MODOMICS were mapped to the positions in the reference tRNA sequences. A local alignment was performed using pairwise2.align.localms from Biopython^40^ with match, mismatch, gap opening and gap extension scores set to 1, −1, −1 and −.1, respectively, and only one alignment reported. Only alignments with a score over 65 were retained. Using the alignments, the modification positions were then lifted over to the fasta sequence positions and the rates of misincorporation and truncations at modified positions were calculated.

### Simulating tRNA-Seq samples

The simulation of tRNA-Seq data was performed using simulateReads.simulate_reads from the simulatetrna python library to generate single end fastq files. Simulated reads contained the ground truth tRNA sequence name in their fastq sequence identifier. Three sets of simulated data were generated.

1. **No modifications:** To compare aligner parameterisation options in the absence of modification-induced truncations and misincorporations, 1 million full length reads were simulated with a sequence error rate of 0.001. One tRNA-Seq sample was simulated from each of the Homo sapiens and Mus musculus tRNA sequences fasta files, with the same number of reads per tRNA.
2. **Uniform:** To compare alignment rates and mis-assignments with reads that approximated the misincorporation profile of real tRNA-Seq data, an equal number of reads was simulated from each tRNA, with a sequencing error rate of 0.01 and truncations and misincorporations added as observed in the error profile. Anticodon-level error profiles were used to simulate truncation and misincorporations by adding these events at a probability equal to the observed frequency. tRNA sequence positions with total misincorporation frequencies less than 0.1 were excluded. One tRNA-Seq sample was simulated from each of the sample-specific error profiles.
3. **Realistic:** To compare the tRNA-Seq quantification approaches, the same approach was taken as for *Uniform*, but with a random number of reads from each tRNA. The random sampling for the number of reads per tRNA was performed using simulateReads.make_gt from the simulatetrna python library. The number of reads was sampled from a left-censured, log2-Gaussian distribution with a mean of 10 and standard deviation of 5. The exponentiated read number was rounded to an integer and values below zero were replaced with zero. A total of 1 million reads were simulated, with the numbers of reads per tRNA sequence down-scaled to achieve this. Ten tRNA-Seq samples were simulated from each of the sample-specific error profiles to enable sample-level evaluation metrics to be obtained.

### Comparing read alignments strategies

*Uniform* simulated samples were aligned to the reference tRNA sequences with bowtie2^22^, SHRiMP^25^. bowtie2 was run with the following non-default parameters: --min-score G,1,8 –local -a -D 20 -R 3 -N 1 -L 10 -i S,1,0.5, where -a makes bowtie2 report all read alignments. SHRiMP was run with the following non-default parameters: --strata --report 1000 --sam-unaligned --mode mirna, where report 1000 makes SHRiMP report up to 1000 alignments per read. The bam.filter_sam function from simulatetrna was used to filter read alignment BAM files from bowtie2 and SHRiMP to remove alignments whose alignment score was less than the highest alignment score for the read. In addition, the mimseq pipeline, which uses GSNAP to align to tRNA sequence clusters was also used. As recommended by mimseq developers (https://github.com/nedialkova-lab/mim-tRNAseq/blob/master/README.md), mimseq was run with the following parameterisation: --cluster-id 0.97--min-cov 0.0005 --max-mismatches 0.075 --max-multi 4 --remap --remap-mismatches 0.05. To compare the read alignments to the ground truths, reads which were multi-mapped by bowtie2 or SHRiMP were assigned in an equal proportion to all aligned coordinates. Ground truth vs assignments for bowtie2 and SHRiMP were compared at Gene locus ID, Transcript ID and anticodon level, where a correct match means that the ground truth and alignment have the same e.g anticodon. The comparison was also made at the mimseq isodecoder level, where the output of mimseq was used to define the mimseq isodecoders. For multi-mapped reads, the match score for multi-mapped reads are bounded between 0 (no alignments are correct) and 1 (all alignments are correct). Ground truth vs assignments for mimseq alignments were compared at anticodon and mimseq isodecoder level.

### Optimising bowtie2 alignment parameters

Bowtie2 was used to align the reads from the *No modifications* simulation samples to the reference tRNA sequences with the following non-default parameters --min-score G,1,8 –local -R 3 -i S,1,0.5 with the value for -D varied from 10 to 100 in steps of 10, the value of -L varied from 10 to 20 in steps of 2, and the value of -N varied from 0 to 1. All combinations of -D, -L and -N were tested. Read alignments were compared to the ground truth as described above.

### Evaluating accuracy of read tallying approaches

*Realistic* simulated samples were aligned to the reference tRNA sequences with bowtie2, SHRiMP, using the same parameters as for the alignment of *Uniform* simulated samples. In addition, the mimseq pipeline, which uses GSNAP to align to tRNA sequence clusters was also used, with the same parameterisation as for the alignment of *Uniform* simulated samples. Five approaches were used to tally reads from the read alignments at the tRNA sequences level (Gene locus ID): *Random, Fractional, Unique, MAPQ>10* and *Salmon*. The read tallying approaches involved the following. *Fractional* - Assign reads in equal fractions to all tRNA sequences to which they are aligned. *Random* - Assign reads at random to one of the tRNA sequences to which they are aligned. *Unique* - Remove all reads aligned to multiple tRNA sequences. Assign remaining reads to their aligned tRNA sequence. *MAPQ>10* - Remove all reads with a mapping quality (MAPQ) score less than 10. MAPQ>10 read tallying was not possible from SHRiMP alignments as SHRiMP does not report a MAPQ. *Salmon* - Use salmon^30^ quant command to estimate the read counts for each tRNA sequence from the alignments.

Read counts were then summed to the transcript ID, anticodon and mimseq isodecoder level, where the output of mimseq was used to define the mimseq isodecoders. *Decision* - assign reads at the Gene locus ID, transcript ID, anticodon and mimseq isodecoder levels. This involved taking the set of alignments for each read and assigning the read at each level where the read was aligned to just a single feature. *Mimseq* - use the mimseq pipeline^16^ to tally reads for each anticodon and isodecoder, where the tRNA sequences within each isodecoder group are defined by mimseq at runtime.

Read tallies per tRNA were compared to the ground truth using two evaluation metrics which were calculated for each tRNA separately: the root mean square error (RMSE)^31^ and the Pearson Correlation coefficient. The evaluation metrics were calculated at Gene locus ID, Transcript ID, anticodon and mimseq isodecoder levels. To compare between quantification approaches, evaluation metrics were further summarised across all features at each level for each tRNA-Seq method by taking the mean value.

### Estimating the contribution of systematic biases to fold-changes

The read alignment and tallying approaches outlined above were applied to the *Uniform* simulated samples and real tRNA-Seq data to quantify the tRNA read counts. For each tRNA-Seq method, fold-changes between pairs of cell-lines/tissues were then calculated at each quantification level for both the *Uniform* simulated sample and real tRNA-Seq samples. The real tRNA-Seq sample log2-fold changes were modeled as being dependent upon the *Uniform* simulated sample log2-fold changes using a least squares linear model with a zero intercept. A separate linear model was used for each level of quantification. The fraction of variance in the real log2-fold changes explained by the log2-fold changes for the simulated sample was obtained from the linear model R-squared value.

## Supplementary Figures

**Supplementary Figure 1.**
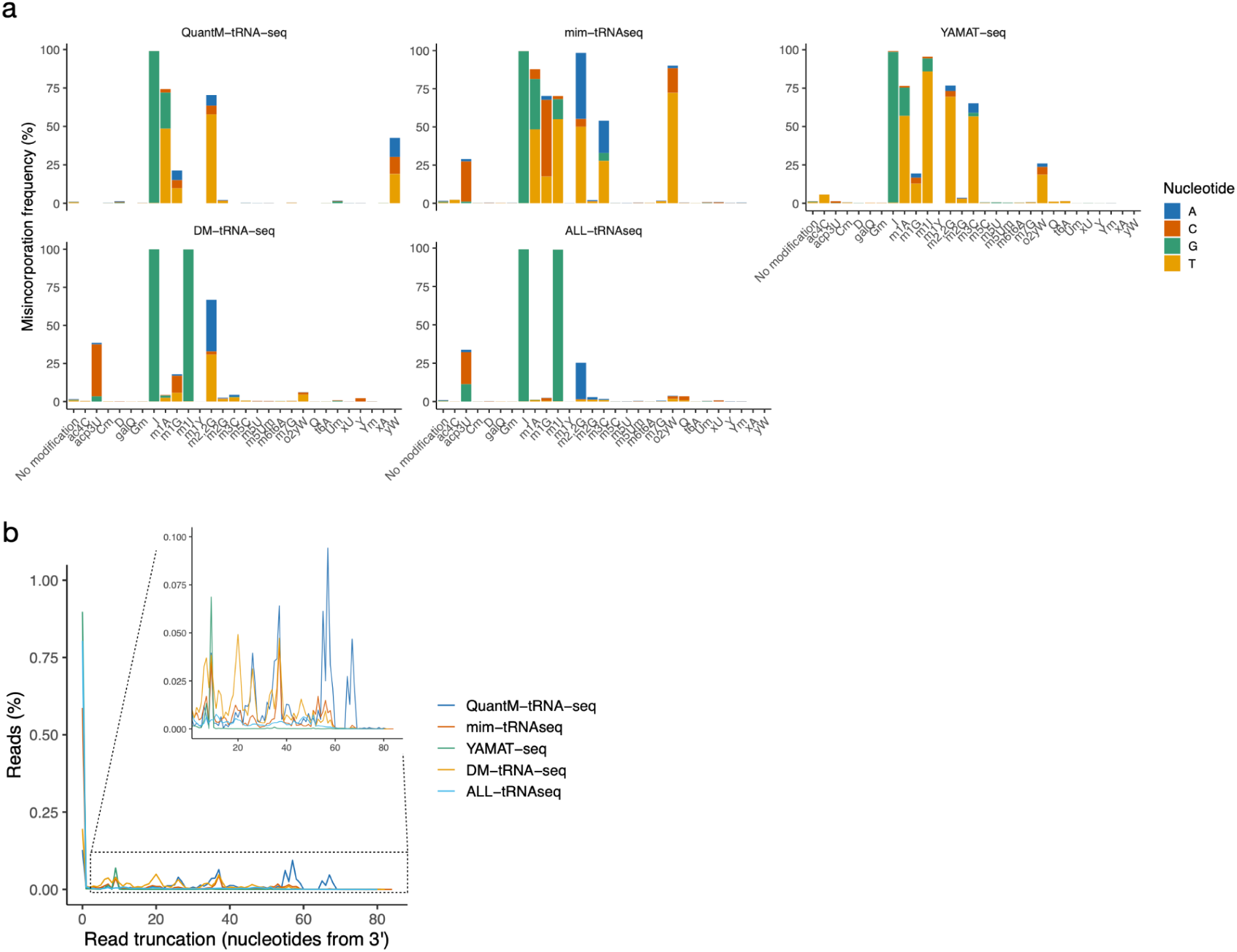
**a)** The complete misincorporation frequencies for all modifications. **b)** The frequency of read truncations with respect to the distance from the 3’ end of the tRNA.

**Supplementary figure 2.**
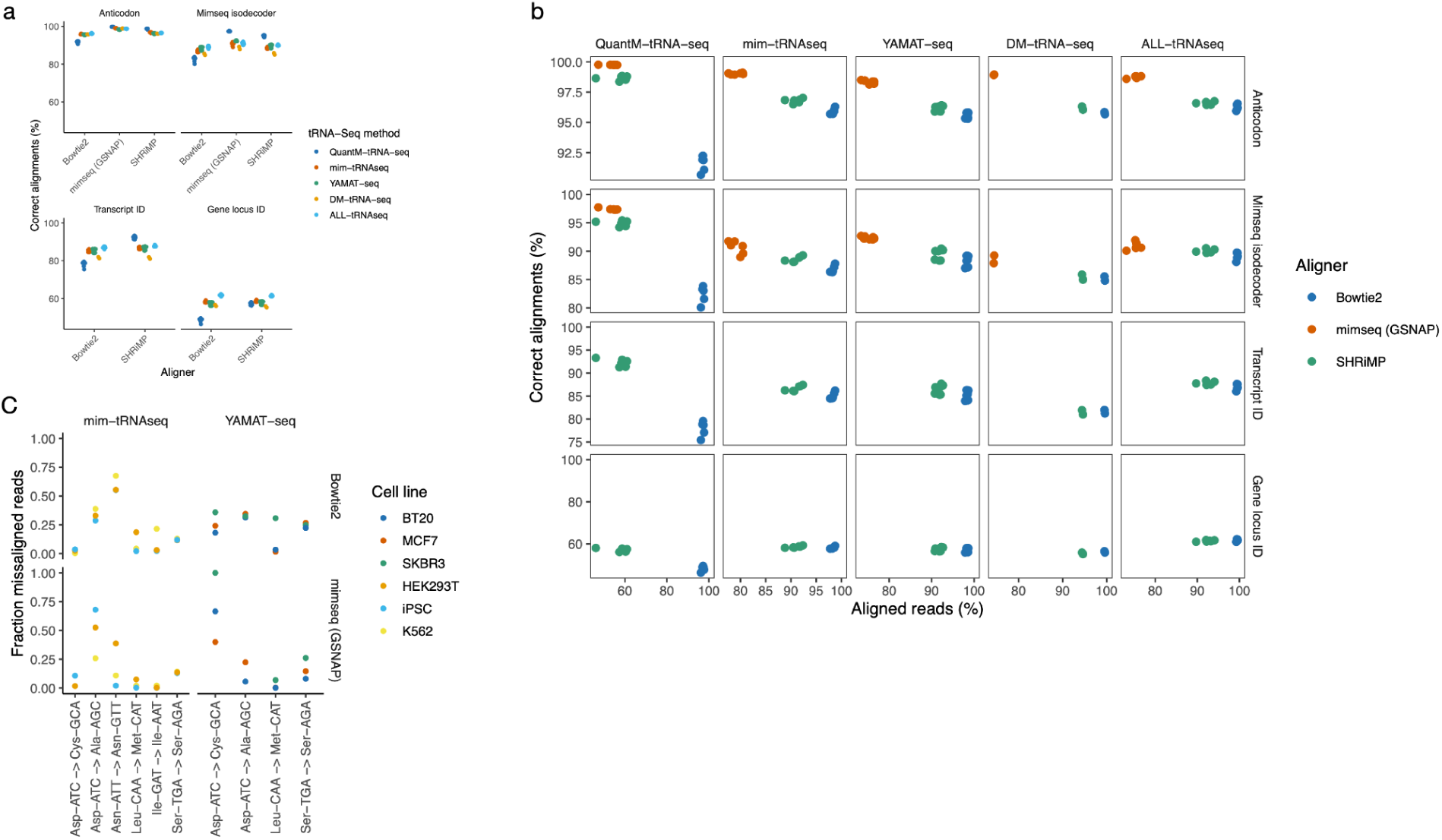
**a)** Percentage of correct alignments at all levels of quantification. **b)** Aligned reads vs correct alignments. Aligners with a higher percentage of aligned reads tend to have a lower percentage of correct alignments. **c)** Example misincorporation events with differential frequency across cell lines in the mim-tRNAseq and YAMATseq simulations.

**Supplementary Figure 3.**
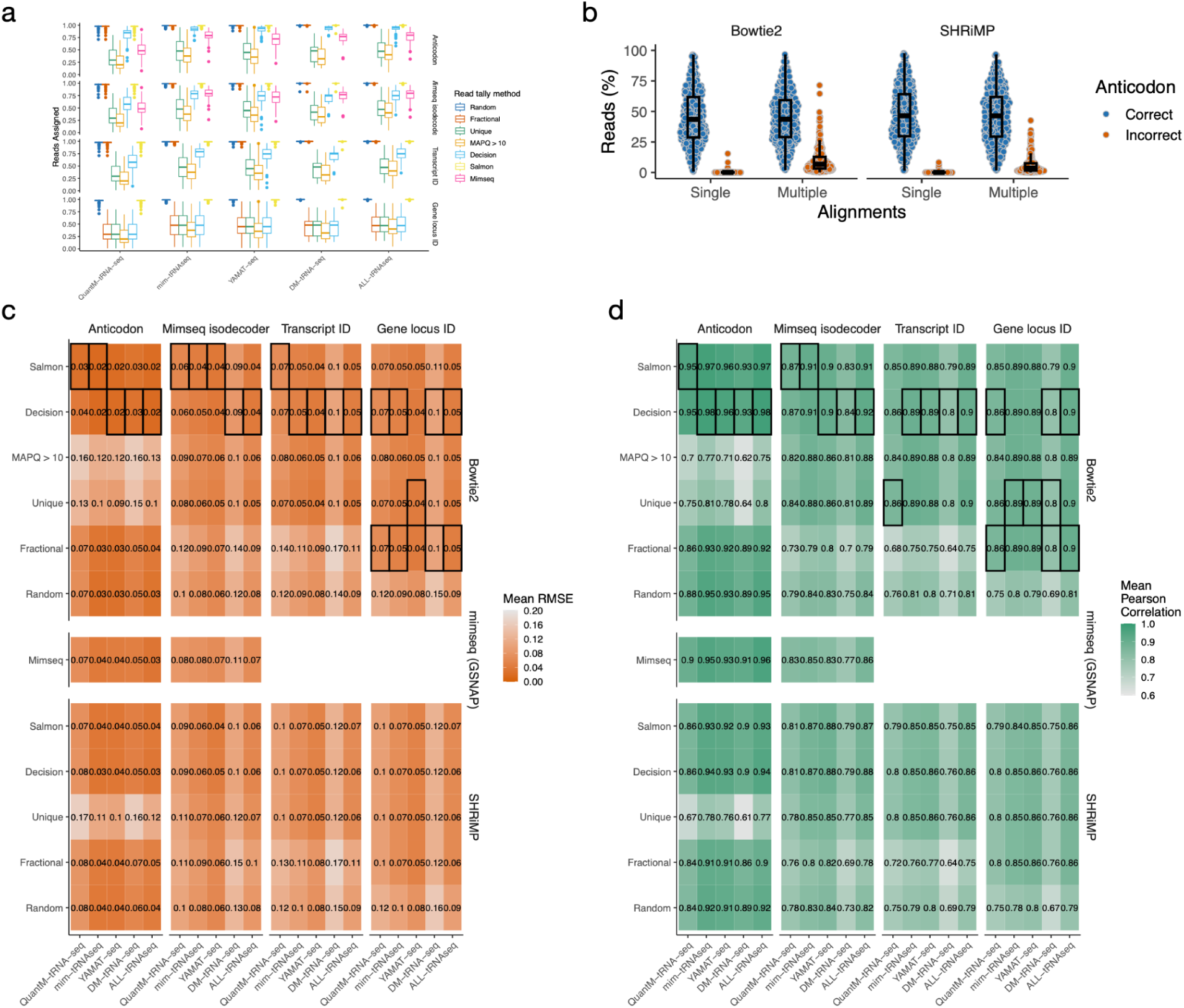
**a** Reads assigned at each level for each read tally method. Only results from alignment with bowtie2 or GSNAP (Mimseq) are shown. **b)** The percentage of reads from each simulation sample which are single or multiple aligned, separated by whether the alignment position(s) have the correct anticodon. Multi-aligned reads with more than more anticodon were deemed incorrect. **c)** Mean RMSE for all approaches at all levels. *=Best approach for each tRNA-Seq method, ▾=Worst approach. **c)** As per (c), except Pearson correlation coefficient.

**Supplementary Figure 4.**
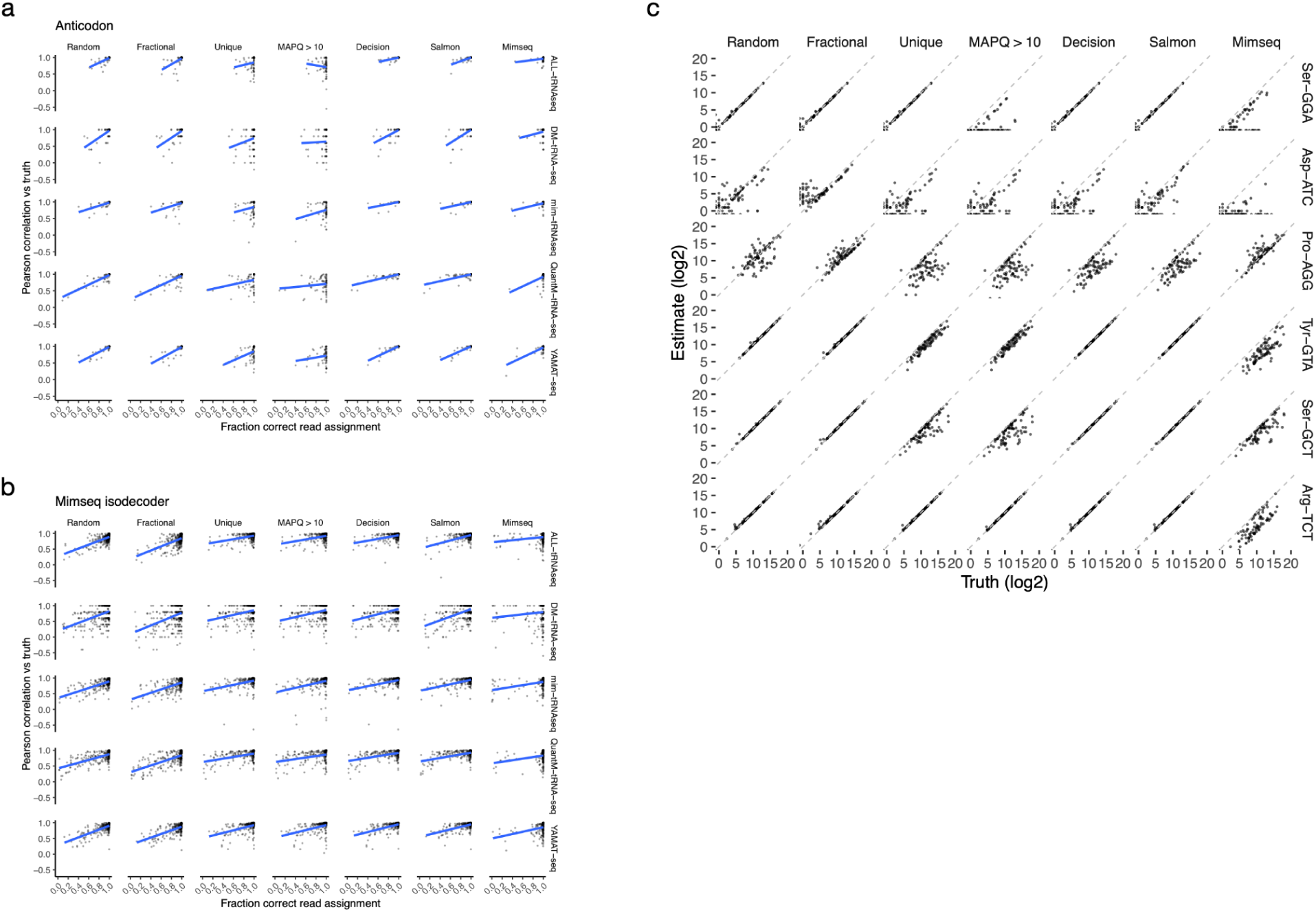
**a).** The correlation between the fraction of reads correctly assigned for each anticodon and the Pearson correlation between the estimated and true read counts. The blue line shows a linear regression fit. **b).** As per (a) but for mimseq isodecoder level quantification. **c)** The correlation between ground truth and estimated read counts for 6 anticodons with variable quantification accuracy, from YAMAT-Seq simulations. The dashed line represents equality.

